# Glucose facilitates Aβ oligomerization and tau phosphorylation in *C. elegans* model of Alzheimer’s disease

**DOI:** 10.1101/228437

**Authors:** Waqar Ahmad

## Abstract

Formation of Aβ plaques from peptide oligomers and development of neurofibrillary tangles from hyperphosphorylated tau are hallmarks of Alzheimer’s disease (AD). These markers of AD severity are further associated with impaired glucose metabolism. However, the exact role of glucose metabolism on disease progression has not been elucidated. In this study, the effects of glucose on Aβ and tau-mediated toxicity are investigated using a *C. elegans* model system. We find that addition of glucose or 2-deoxy-d-glucose (2DOG) to the growth medium delayed Aβ-associated paralysis, though it was unable to restore previously impaired acetylcholine neurotransmission in pre-existing Aβ-mediated pathology. Glucose also inhibited egg laying and hatching in the worms that express Aβ. The harmful effects of glucose were associated with an increase in toxic Aβ oligomers. Increased phosphorylation of tau is associated with formation of neurofibrillary tangles (NFTs) and increased severity of AD, but O-β-GlcNAcylation can inhibit phosphorylation of adjacent phosphorylation sites. We reasoned that high glucose levels might induce tau O-β-GlcNAcylation, thereby protecting against tau phosphorylation. Contrary to our expectation, glucose increased tau phosphorylation but not O-β-GlcNAcylation. Increasing O-β-GlcNAcylation, either with Thiamet-G (TMG) or by suppressing the O-GlcNAcase (*oga-1*) gene does interfere with and therefore reduce tau phosphorylation. Furthermore, reducing O-β-GlcNAcylation by suppressing O-GlcNAc transferase (*ogt-1*) gene causes an increase in tau phosphorylation. These results suggest that protective O-β-GlcNAcylation is not induced by glucose. Instead, as with vertebrates, we demonstrate that high levels of glucose exacerbate disease progression by promoting Aβ aggregation and tau hyperphosphorylation, resulting in disease symptoms of increased severity. The effects of glucose cannot be effectively managed by manipulating O-β-GlcNAcylation in the tau models of AD in *C. elegans.* Our observations suggest that glucose enrichment is unlikely to be an appropriate therapy to minimize AD progression.

## Introduction

Alzheimer's disease (AD), an age-associated, irrevocable brain disorder, results in chronic neurodegeneration and is the most common cause of dementia. The accumulation of extracellular Aβ plaques and hyperphosphorylated tau neurofibrillary tangles (NFTs) in the AD brain has been identified as a major cause of toxicity (1-4).

Most AD therapies focus on decreasing the Aβ levels and tau hyperphosphorylation. Hyperphosphorylation of tau leads to self-aggregation into tangles of filaments that cause a loss of neuronal function and neuronal degeneration (5, 6). Tau O-β-GlcNAcylation is reciprocally antagonistic to phosphorylation and has been proposed as a possible therapy to reduce tau phosphorylation (5, 7, 8). Glucose metabolism through the hexosamine biosynthesis pathway was proposed to induce O-β-GlcNAcylation, whereas a decrease in intracellular glucose levels were suggested to be associated with reduced O-β-GlcNAcylation and increased tau phosphorylation in AD brains (9, 10). However, no study has directly tested the effect of elevated glucose levels on tau O-β-GlcNAcylation and phosphorylation.

An opposing model of the interaction between glucose and AD involves energy metabolism. In one iteration, this energy metabolism model suggests that impaired glucose metabolism causes AD. This is supported by the observation that reduced glucose metabolism and associated enzyme activities were observed in AD patients (13-18). In contrast, there is a decrease in Aβ-toxicity after caloric restriction or inhibition of glucose catabolism, which indicates that reduction in glucose-associated metabolism protects against neurodegeneration (19-25). These contradictory observations mark glucose metabolism as an important target for studies designed to increase our understanding of the progression of AD.

In this study, we have tested the outcome of glucose enrichment in the AD model of the simple invertebrate *C. elegans* either expressing Aβ or tau. A drug known as Thiamet G (TMG) that induces O-β-GlcNAcylation was also used in our study along with glucose to examine the effect of O-β-GlcNAcylation on Aβ toxicity and tau phosphorylation.

## Materials and methods

### Nematode strains

Wild-type strain, N2 (Bristol), human β-amyloid expressing strains CL2006 (dvIs2 [pCL12(unc-54/human Abeta42 minigene) + pRF4]), CL4176 (dvIs27 [(pAF29)myo-3p::A-Beta (42)::let 3’UTR) + (pRF4)rol-6(su1006)]), CL2355 (dvIs50 [pCL45 (snb-1::Abeta 1-42::3’ UTR(long) + mtl-2::GFP] I), and human tau expressing strain VH255 (hdEx82 [F25B3.3::tau352(WT) + pha-1(+)]) was used for this study. No Aβ-control for CL2355 was CL2122 (dvIs50 [(pD30.38) unc-54 (vector) + (pCL26) mtl-2::GFP] I). In muscles, CL2006 produces the human Aβ peptide constitutively while CL4176 expresses the Aβ peptide when the temperature is increased from 16°C to 23°C in muscles. CL2355 express Aβ in neuronal cells. The use of these strains as a worm model of AD was documented previously (26). The strain VH255 expresses fetal 352aa CNS tau in neurons (27). Strain RB1342 [*ogt-1*(ok1474)III] was used as –ve control for O-GlcNAcylation study.

### Culture conditions

The cultures of *C. elegans* were maintained on nematode growth medium (NGM) seeded with *E. coli* OP50 at 20°C, except strains CL4176 and CL2355 that were maintained at 16°C to suppress Aβ expression. Synchronized cultures of nematodes for bioassays were obtained by harvesting eggs from gravid hermaphrodites by exposing them to a freshly prepared alkaline bleach solution (0.75N NaOH + 1.5N NaOCl). The worms were incubated in the bleach solution for five minutes at room temperature followed by centrifugation at 3300 RPM for 1 minute at room temperature. The supernatant was then discarded, and the pelleted eggs were resuspended in M9 buffer (6 g/L Na_2_HPO_4_; 3 g/L KH_2_PO_4_; 5 g/L NaCl; 0.25g/L MgSO_4_ •7H_2_O. Washing with M9 buffer was repeated 3 more times, after which the eggs were suspended in M9 buffer and allowed to hatch overnight on an orbital shaker. The resulting L1 larvae were shifted to fresh NGM agar plates seeded with *E. coli* to initiate growth. To select array in VH255 strain, eggs were shifted to freshly made NGM plates and placed at 25°C overnight.

Aβ transgenic worms were initially cultured at 16°C for 36 hours, after which the temperature was increased to 23°C for 36 hours, except for the paralysis assay, for which the temperature was further increased to 25°C to maximize expression of the Aβ transgene. For VH255, worms were propagated at 20°C after hatching for further analysis. The phenotypes of the worms were monitored by visual observation under a microscope and/or quantified using the WormScan procedure (28).

### Oga-1 and ogt-1 gene suppression

The *E. coli* strains SJJ_T20B5.3 and SJJ_K04G7.3 (Source Bioscience), which expresses double-stranded RNA of the *oga-1 and ogt-1* genes, respectively, were fed to *C. elegans* strains to suppress the expression of the *dld-1*, *oga-1 and ogt-1* genes, respectively (29). *E. coli* strain HT115 containing empty vector was used as a control in these experiments. Briefly, the bacteria were cultured in LB medium containing 100μg/ mL ampicillin overnight with shaking at 37°C. The 300 μL of this bacterial culture was transferred to NGM plates containing 100μg/mL ampicillin and 1mM isopropyl β-D-1-thiogalactopyranoside (IPTG). The plates were incubated at 25°C overnight to allow the bacteria to grow. Synchronized L1 worms were transferred to the bacterial plates and kept at 16°C for 36 hours. After a further 36 hours at 23°C, the worms were ready for use in the assays described below.

### Paralysis assay

Synchronized L1 stage worms of strain CL4176 that express Aβ were used in paralysis assay. To assess the role of glucose or glycerol on paralysis behaviour of Aβ transgenic strains, distinctive concentrations (0.5%, 1%, 2%, 4%, and 10%) were used in normal NGM plates. For TMG, distinct concentrations ranging from 2μM-80 μM was used. The effect of 5mM 2-deoxy-d-glucose (2DOG) on worms expressing Aβ was also studied. The control strain for these assays was CL4176 worms fed with OP50. For RNAi experiments, worms transferred to NGM plates that had been seeded with the *oga-1 or ogt-1* ds-RNA *E. coli* strains to achieve gene suppression or with the *E. coli* strain HT115 containing the empty vector as a control. After 36 hours at 16°C, worms that express Aβ were up shifted to 25°C and scored for paralysis every 24 hours after an initial 24-hour period until the all worms became paralyzed.

### Aldicarb and levamisole assays

L1 stage worms were incubated with different glucose concentrations at 20°C after synchronization. After 72 hrs worms were transferred to the plates either with 1mM aldicarb (an acetylcholinesterase inhibitor (30)) or 0.2mM levamisole (a cholinergic agonist (31)). The number of active worms was counted every half an hour until all worms become paralyzed.

### Fecundity and egg hatching assay

Worms were synchronized and grown on NGM agar seeded with OP50 bacteria at 16°C. Once they reached maturity at four days of age, 10 individuals in each of 3 replicate were transferred to fresh agar plates of the same composition and shifted to 20°C. Unhatched eggs and larvae were counted every 24 hours for the next three days.

### Liquid thrashing assay

Liquid thrashing assays were performed using at least 10 worms from each cohort every 2^nd^ day until day 7. Thrashes were counted in a time interval of 10 seconds at 20°C.

### Glucose measurement

Synchronized day 3 (L4) worms were washed and 400μL ice cold RIPA buffer was added to each sample. Worms were sonicated, and the supernatant was collected in new tubes. Soluble protein was quantified and used as a normalizing factor during glucose quantification. This supernatant was subjected to glucose quantification by Sigma glucose assay kit (GAG*O*-20) following the manufacturer guidelines.

### Measurement of worm’s speed

Wild type worms were synchronized and placed on normal NGM plates or plates supplemented with glucose at concentrations ranging from 0.5%-10% seeded with OP50, and placed at 20°C. Speed of movement was measured using WormLab 4.0 software (http://www.mbfbioscience.com/wormlab). Worm movements were tracked for 3 days.

### Western Blotting

Aβ was identified in *C. elegans* strains by immunoblotting after separation on a 16% Tris-Tricine gel. A standard western blotting protocol was used except that SDS was omitted from the transfer buffer. Briefly, synchronized L4 worms were incubated at 23°C for 48h, then washed with distilled water and quickly frozen in liquid nitrogen. Flash frozen worms were either stored at -80°C or sonicated twice in the ice-cold cell lysis buffer (50 mM HEPES, pH 7.5, 6 mM MgCl_2_, 1mM EDTA, 75 mM sucrose, 25 mM benzamide, 1 mM DTT and 1% Triton X-100 with proteinases and phosphatases inhibitors 1:100 ratio). After sonication, the lysate was centrifuged at 10000 rpm to remove insoluble debris and soluble protein in the supernatant was measured using Pierce Coomassie (Bradford) protein assay kit (Thermo Scientific) on Nanodrop. From each sample, 80-100 μg of total protein was precipitated with acetone and dissolved in Novex^®^ Tricine SDS sample buffer (LC1676, Invitrogen) by heating to 99°C for 5 minutes. Aβ samples were subjected to gel electrophoresis at 100V for 2.5 hrs in the separate cathode (100mM Tris, 100mM Tricine, 0.1% SDS, pH 8.3) and anode (0.2M Tris, pH 8.8) running buffers. Proteins were semi-dry transferred onto nitrocellulose membranes by electroblotting in the transfer buffer (35 mM glycine, 48 mM Tris (pH = 8.8) and 20% methanol) for 60 min at 300mA and stained with Ponceau S (0.1% Ponceau S in 1% acetic acid) for 5 minutes following de-staining with 10% acetic acid (5 minutes) and washing under water to completely remove the acetic acid (5 minutes).

For tau, synchronized worms were incubated at 20°C and L4 worms (day 3) were collected and washed three-four times to completely remove the bacterial traces. Half of the washed worms were added in 400μl cold RIPA buffer containing proteinases and phosphatases inhibitors. The remaining half of the worms were shifted to new plates containing 75μM FUDR to restrict progeny production and collected on day 7. Worms were subjected to sonication, and lysate was collected for further protein analysis. Protein was measured and about 50-80μg protein was subjected to 10% SDS-PAGE gels for two hours. Proteins were transferred to nitrocellulose membranes for further analysis in SDS-containing transfer buffer.

For Aβ, the membranes were blocked overnight in 5% skim milk at 4°C to prevent nonspecific binding of antibodies. For tau protein extracts; membranes were blocked in 5% BSA in 0.1% TBST ((TBS containing 0.1% Tween 20). Primary antibody staining was achieved using the Aβ monoclonal antibody 6E10 (Covance) at 1:1000 dilution in TBS (50mM Tris, 150mM NaCl, pH 7.6) containing 1% skim milk for 3-4 hours at room temperature following three washes with TBS-T five minutes each. A number of antibodies were used to assess total tau and site-specific phosphorylated tau. The antibodies used for tau were, Tau-5 (ab80579, Abcam), HT7 (MN1000, Thermo Scientific), phosphor S198 (ab79540, Abcam), phosphor S235 (NB100-82241, Novus Biologicals), phosphor S262 (79-152, Prosci), phosphor S396 (710298, Novex Lifetecnologies), AT8 (S202/ Thr-205, MN1020, Thermo Scientific), and AT180 (T231/ Ser-235, MN1040, Thermo Scientific). For O-GlcNAc detection, anti-O-GlcNAc antibody (CTD110.6, sc-59623, SantaCruz) was used. Anti-actin antibody (2Q1055, Abcam) or anti-γ-Tubulin (T1450, Sigma) was used as reference control. Anti-mouse IgG alkaline phosphatase antibody produced in goat (A3562, Sigma), and anti-rabbit IgG alkaline phosphatase antibody produced in goat (A3687, Sigma was used as secondary antibody at 1:10000 dilution in TBS containing 1% skim milk or 1% BSA in 1%TBST. Secondary antibody staining was done for one hour at room temperature. After washing the membrane with 1X TBS-T, the proteins were detected using BCIP/ NBT substrate system (Sigma) or BCIP/ NBT kit (002209) from Lifetechnologies dissolved in 1M Tris (pH 9.0).

### Statistical analysis

Differences due to treatments, strains and epigenetic gene suppression were analyzed for statistical significance by independent student’s *t*-tests for two groups and one-way ANOVA for multi-groups using GraphPad prism 0d. Log-rank estimate was applied in survival calculations. A *p* value less than 0.05 was considered statistically significant.

## Results

Glucose is an essential source of energy generation in neurons whereas a decline in glucose metabolism was evident in patients with AD. Here in this study, we assessed any possible link between Aβ and tau-mediated toxicity and glucose enrichment using transgenic *C. elegans* strains that express the human peptides.

### Glucose, glycerol, and 2DOG delay paralysis caused by Aβexpression in C. elegans muscle

Production and accumulation of Aβ in the AD brain is an important hallmark in disease progression. Whereas, accretion of Aβ in human muscles resulted in reduced functioning (32, 33). Temperature inducible expression of muscle Aβ in *C. elegans* resulted in a time-dependent paralysis phenotype. (34). Our results showed that supplementation of the growth medium with glucose increases intracellular glucose levels (Fig S1) and delays paralysis in a dose-dependent manner. The untreated worms that express Aβ were completely paralyzed within 48 hours with a median time of 24 h. Though addition of 0.5% glucose significantly delayed paralysis (p<0.0001, median time=48h, max survival time=72h), the maximum delay in paralysis occurred at 1% glucose (p<0.0001, median time=72h, max survival time= 144h). Higher concentrations of glucose increasingly reversed this delay (2%glucose: p<0.0001, median time=60h, max survival time= 144h; 4%glucose: p<0.0001, median time=48h, max survival time= 96h; 10%glucose: p=0.059, median time=48h, max survival time= 72h).

To determine whether the decrease in paralysis due to glucose (Fig 1A) is glycolysis dependent, we treated the worms with 5mM 2DOG, an enantiomer of D-glucose. D-glucose cannot be phosphorylated by hexokinase and is thus unable to enter the glycolytic pathway, creating a dietary restriction-like state. 5mM 2DOG significantly reduced paralysis in worms that express Aβ (72 hrs vs 96 hrs, p<0.0001, median survival= 48h) (Fig 1B). Our results showed glycolysis-independent mechanism of glucose-mediated protection against Aβ-proteotoxicity.

**Figure 1:**
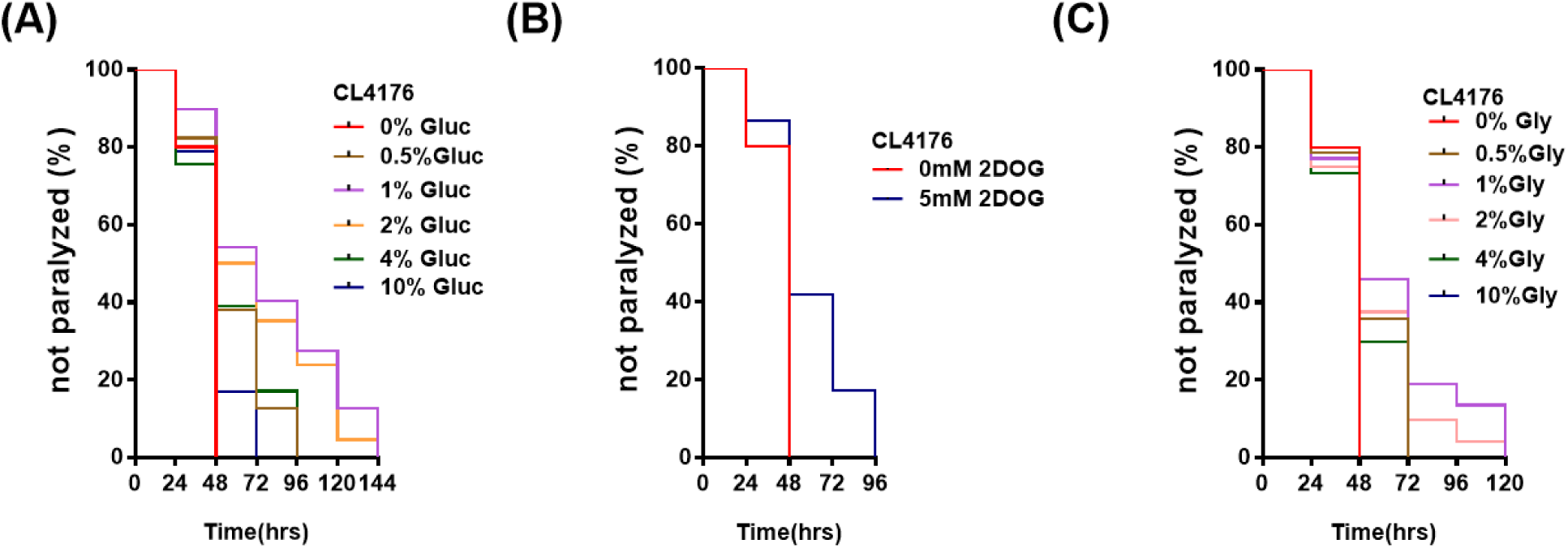
Glucose, 2DOG, and glycerol alleviates paralysis due to expression of Aβ in muscles. Standard NGM plates seeded with *E. coli* OP50 were prepared containing different concentrations of glucose or glycerol, or 5mM 2DOG. Worms were assessed for paralysis at 24 hrs interval after temperature upshift. Three independent trials were run for each group, and results were presented as the weighted average. Log-rank test was used to determine any significant change in paralysis behavior between each group. N=50-60 worms per replicate for each data point. (A) Effect of increasing glucose on paralysis of CL4176 strain. (B) Effect of 5mM 2DOG on paralysis. (C) Effect of increasing glycerol on Aβ-mediated paralysis.

Glycerol can also be used as a carbohydrate energy source and contribute to oxidative phosphorylation (35, 36). Similar findings as of glucose were observed in worms that express Aβ when fed with glycerol. At 0.5%, 1%, and 2% glycerol in NGM medium, decreased paralysis was observed. Maximum relief against paralysis was observed when worms were fed with 1% glycerol (48 hrs vs 120 hrs, p<0.0001, median time= 48h). At higher concentration of glycerol, the paralysis rate began to rise (48 hrs vs 72 hrs, p=0.026, median time= 48h) with all worms dying at 10% glycerol within 24 hours. (Fig 1C).

### Glucose impairs cholinergic neurotransmission in muscle of C. elegans

Expression of Aβ in *C. elegans* muscles is associated with impaired ACh neurotransmission (37). Given that glucose protects against Aβ-mediated paralysis in *C. elegans*, we were interested to determine whether this improvement was due to improved ACh signaling. Moreover, Aβ expression in *C. elegans* blocks ACh release, any improvement after glucose of 2DOG administration will also effect the Aβ physiological state (38). The strain CL2006 was used to assess the effect of glucose or 2DOG on pre-existing Aβ pathology while, wild type worms were used as control. The median paralysis time for wild type was 30 min while worms that express Aβ have induced median survival time to 60 min either treated with aldicarb (an ACh esterase inhibitor) or levamisole (a cholinergic agonist). Glucose supplementation of the medium did not restore normal ACh neurotransmission in CL2006, but rather seemed to further impair cholinergic signaling as it increased resistant to both aldicarb and levamisole. Similar results were observed in wild type worms (Fig 2, Table S1), indicating that the effect was independent of Aβ.

**Figure 2:**
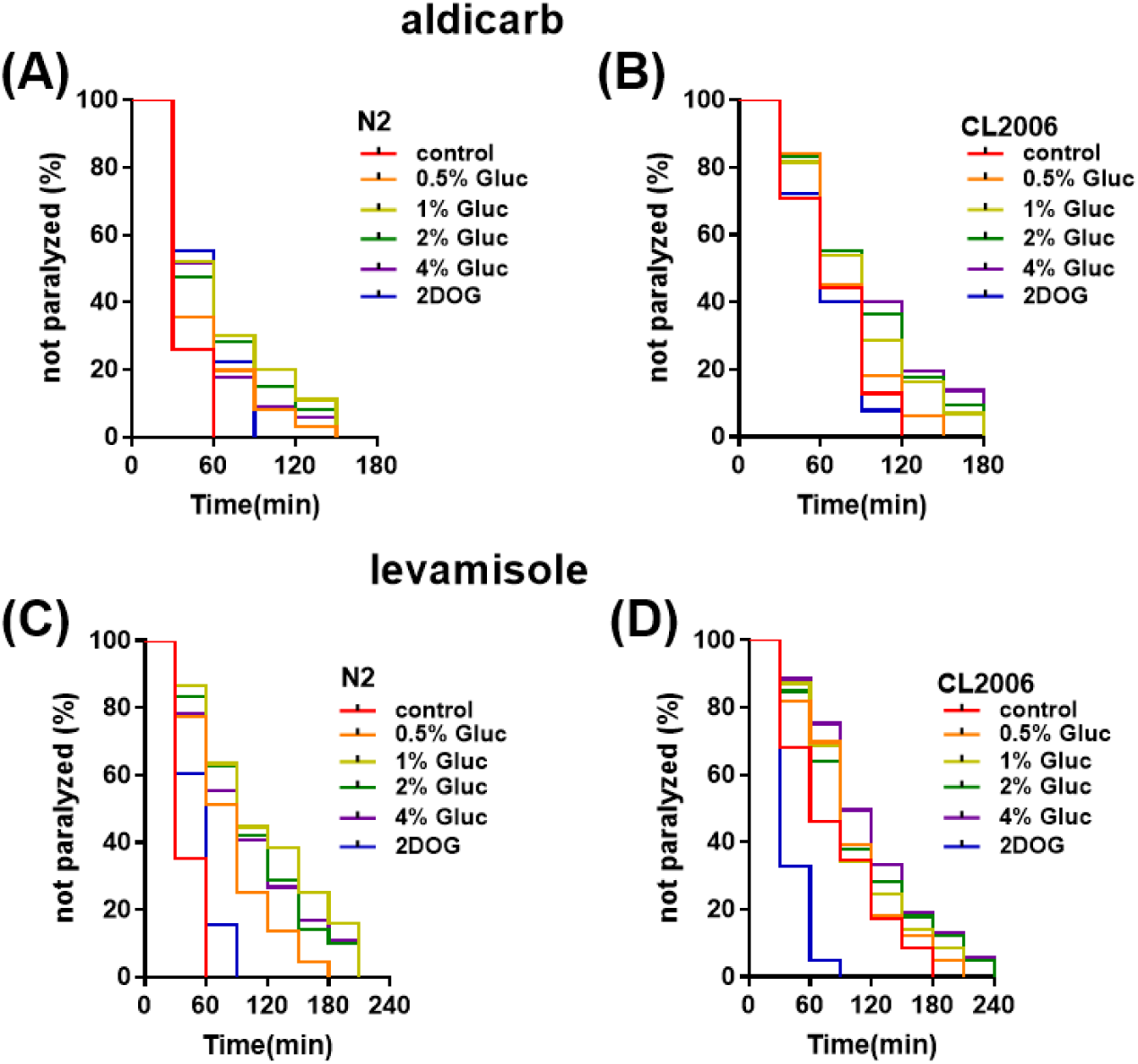
Effect of glucose and 2DOG on ACh-mediated neurotransmission. Addition of glucose impaired while 2DOG restores the ACh neurotransmission in muscles of worms that express Aβ. Wild type and Aβ constitutively expressing strain CL2006, were exposed to different concentrations of glucose or 5mM 2DOG for 72h and then transferred to freshly prepared NGM plates containing aldicarb or levamisole. Three independent trials were run for each group, and results were presented as the weighted average. Log-rank test was used to determine any significant change in paralysis behavior between treatments (n = ~50 worms per trial). Table 1 shows summary of the statistical analysis. A) Wild type worms in the presence of 1mM aldicarb. B) CL2006 worms in the presence of 1mM aldicarb. C) Wild type worms in the presence of 0.2mM levamisole. D) CL2006 worms in the presence of 0.2mM levamisole.

**Table 1:** Effect of glucose, glycerol, and 2DOG on worm egg laying and hatching

|  |  |  |  |  |  |
| --- | --- | --- | --- | --- | --- |
| **Treatment** | **Total No. of Eggs** | **Unhatched Eggs (%)** |  |  |  |
|  |  | **Day 0** | **Day 1** | **Day 2** | **Day 3** |
| **N2** | 294±34 | 100 | 10.6±1.5 | 1±0.5 | 3.6±0.8 |
| **CL2122** | 273±23 | 100 | 39±7 | 19.6±3.5 | 8.6±2.1 |
| **CL2355** | 157±7 | 100 | 61.8±4.4 | 37.5±5.1 | 17.3±3.4 |
| **CL2355 (0.5%G)** | 164±11 | 100 | 40.1±10.9 | 25.1±3.7 | 13.1±3.1 |
| **CL2355 (1%G)** | 133±17 | 100 | 45.9±3.4 | 27.7±5.1 | 17.1±3.9 |
| **CL2355 (2%G)** | 117±8 | 100 | 45.5±2.3 | 27.4±4.5 | 11±2.1 |
| **CL2355 (4%G)** | 109±16 | 100 | 58.4±3.1 | 42.3±5.1 | 28.5±4.7 |
| **CL2355 (10%G)** | 62±9 | 100 | 62.4±1.7 | 49.4±0.5 | 40.3±5.1 |
| **CL2355 (0.5% GLY)** | 177±14 | 100 | 50.6±8 | 25.9±3.9 | 14.6±3.6 |
| **CL2355 (1% GLY)** | 214±31 | 100 | 44.3±1.3 | 22.5±4.8 | 12.1±1.7 |
| **CL2355 (2% GLY)** | 131±12 | 100 | 28.1±5.1 | 15±4.1 | 8.5±3.3 |
| **CL2355 (4% GLY)** | 81±20 | 100 | 52.3±8.8 | 38.5±3.2 | 30.5±4.6 |
| **CL2355 (2DOG)** | 173±35 | 100 | 61.9±3.3 | 32.6±2.5 | 19.5±4.2 |

In the presence of aldicarb, treatment with 5mM 2DOG had no effect on paralysis in Aβ transgenic worms (p=0.59), but it increased time to paralysis in wild type worms (60min vs 90 min, p<0.0001) (Fig 2A and 2B). In the presence of levamisole, 5mM 2DOG restored ACh neurotransmission in CL2006 (180min vs 90min, p<0.0001) while wild type worms showed resistance against 2DOG (60min vs 90min, p<0.0001 (Fig 2C and 2D).

To evaluate the effect of glucose, glycerol and 2DOG on the Aβ-mediated decrease in fertility, we used strain CL2355 that expresses human Aβ panneuronally along with its matched control CL2122. There was no difference in the number of eggs laid between wild type (N2) and CL2122 worms (294±34 vs 273±23, p>0.05), but the time required for the eggs to hatch was delayed in strain CL2122 (p=0.002). Expression of Aβ reduced fecundity of CL2355 compared to CL2122 (273±23 vs 157±7, p<0.001) and also delayed egg hatching with 61% unhatched eggs (p=0.008) (Table 2).

Supplementation of the growth medium with glucose resulted in a dose dependent decrease in fecundity at concentrations ranging from 1% to 10% in worms that express Aβ (Table 2). Interestingly, we observed a significant increase in fecundity at 1% glycerol (157±7 vs 214±31, p<0.05) in worms that express Aβ. From 2% to 4% glycerol however, there was a dose dependent decrease in fecundity. The apparent increase in the number of eggs laid in response to 2DOG was not statistically significant (157±7 vs 173±31, p= 0.432). Glucose, glycerol and 5mM 2DOG each significantly reduced egg hatchability. Worms fed on >2% glucose or glycerol were most severely affected with more than 30% of eggs remaining unhatched by day 3.

### Glucose and glycerol, but not 2DOG, induce Aβ oligomerization

As our results seemed to indicate that the deleterious effects of glucose were independent of Aβ, we wished to determine whether glucose had any effect on Aβ oligomerization. The production and accumulation of Aβ oligomers are critical to the progression of AD resulting in long-term potentiation and memory impairments (39-41). To investigate Aβ oligomerization, soluble proteins from whole cell lysates of worms grown on glucose, 2DOG, or glycerol were subjected to western blotting. Our results show that higher concentrations of glucose (4% and 10%) decreased the ~4kDa Aβ monomers (1.19 and 2.45-fold, respectively) and substantially increased the ~9kDa oligomers (1.66 and 1.59 fold, respectively) and the ~16kDa oligomers (1.26 and 1.35 fold, respectively) compared to the levels in control worms cultured without glucose. We observed similar results to glucose when worms were fed with different concentrations of glycerol, where a gradual increase in glycerol induced Aβ oligomerization (Fig 3B). Interestingly, 5 mM 2DOG was not found to induce Aβ oligomerization. Rather, it significantly increased ~4kDa monomers (1.56 fold, p = 0.045) with a reduction in ~16 kDa oligomers (1.51 fold, p = 0.048) when compared to untreated control (Fig 3A).

**Figure 3:**
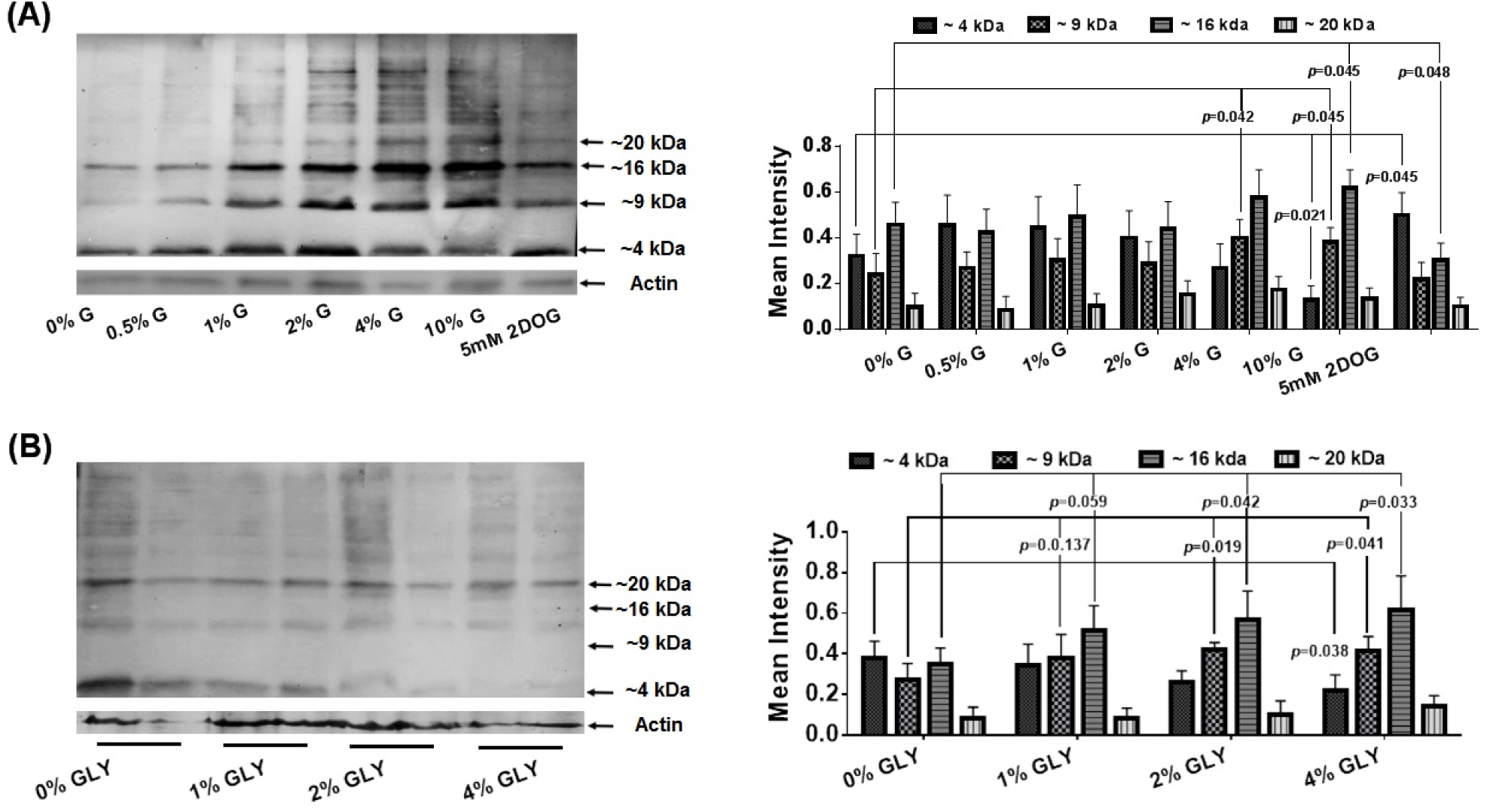
Effect of glucose, 2DOG, and glycerol on Aβ oligomerization. Synchronized L1 stage CL4176 worms were treated with v various concentrations of glucose or glycerol, or 5mM DOG at 16°C for 36 h, followed by 36 h at 23°C to induce Aβ transgene expression. Total cell lysate proteins were subjected to 16% Tris-Tricine SDS-PAGE and detected by anti-Aβ antibody 6E10. Anti-actin antibody was used as reference control. Original uncropped images with protein ladder (ab116029) are provided in supplemental section (Fig S2). Quantification of Aβ monomers and oligomers appeared on gel was performed using GelQuantNET software. Graphs show results from two biologically independent experiments. A) Aβ transgenic worms were fed with different concentrations of glucose (0.5%-10%) or 5mM 2DOG. B) Western blot of Aβ strain CL4176 grown on medium supplemented with various concentrations of glycerol (1%-4%). Error bars = mean ±SD.

### Glucose induces tau phosphorylation

Like Aβ oligomerization, tau hyperphosphorylation is an important contributor to AD pathology as it promotes the formation of neurofibrillary tangles (42). We examined tau phosphorylation in worms grown on 1% or 4% glucose. We observed increased phosphorylation at tau Ser-198 (1%G, 12.9 fold; 4%G, 15.7 fold), Ser-235 (1%G, 12.8 fold; 4%G, 18.4 fold), and Ser-262 (1%G, 5.7 fold; 4%G, 7.1 fold) when compared to control worms grown on medium without glucose supplementation (Fig 4). No change in the overall level of phosphorylation of the tau protein in response to glucose was observed using the HT-7 antibody. This highlights the specificity of the dramatic changes that were observed using the other antibodies.

**Figure 4:**
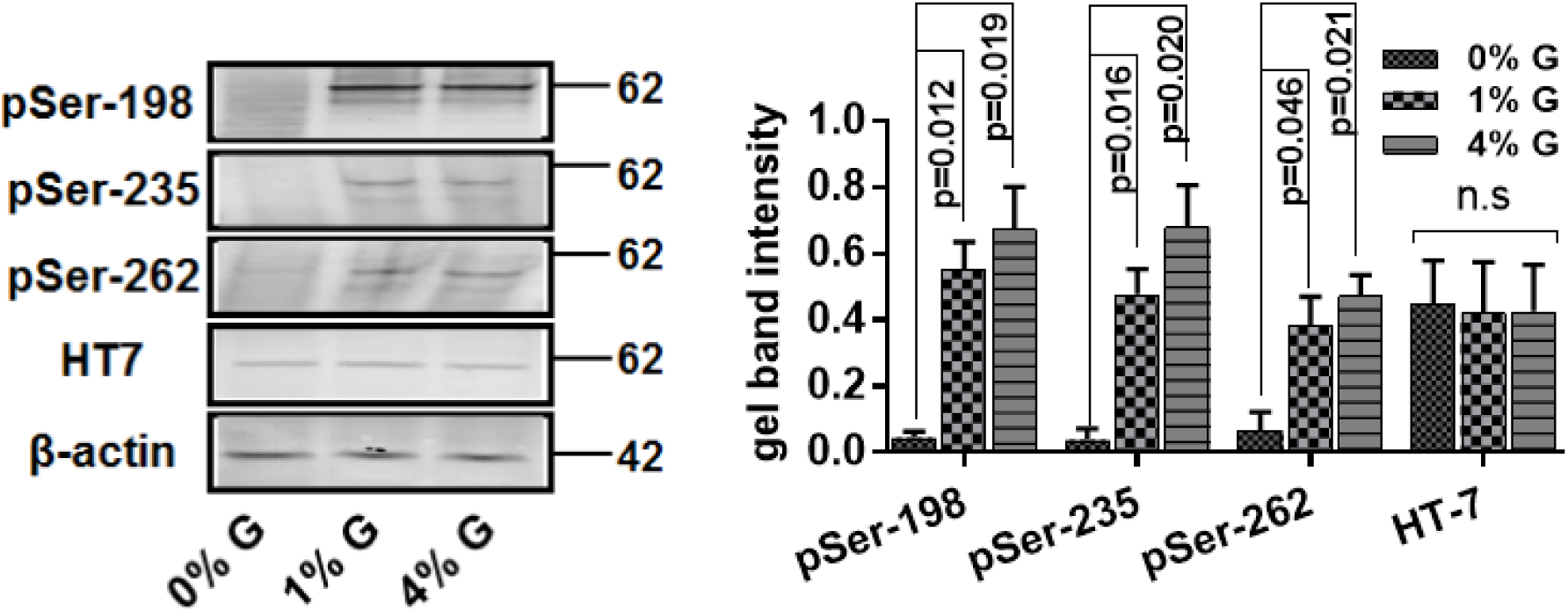
Effect of glucose on site-specific phosphorylation of tau expressed in *C. elegans.* Synchronized L1 stage worms of strain VH255 expressing human fetal tau were treated with 1% or 4% glucose for 72 h at 20°C. Total cell lysate protein was extracted and subjected to western blot. Tau phosphorylation was monitored using different phosphor-tau-antibodies. Total tau was detected using HT7 antibody while, anti-actin antibody was used as reference control. Graphs indicate results from two independent biological trials. Bars= Mean ± S.D.

### Oga-1 suppression modulates Aβ-mediated paralysis in worms that express Aβ in muscle, and tau phosphorylation in human tau-expressing worms

Hyperglycemia has been proposed to induce O-β-GlcNAcylation, which in turn has been proposed to impede phosphorylation of adjacent phosphorylation sites (10). Contrary to expectation, we observed increased phosphorylation of tau in response to increased levels of glucose in the growth medium. To understand the basis of this result, we sought to modulate the level of O-β-GlcNAcylation directly and look at its impact on tau phosphorylation. We used a mutation in the O-GlcNAc transferase gene (*ogt-1*) to prevent O-glycosylation. We also used a mutation in the O-GlcNAcase gene (*oga-1*), as well as thiamet G, an O-GlcNAcase inhibiting drug proposed to treat AD, as both of these prevent O-GlcNAc removal (10, 43-47).

TMG caused a dose dependent delay in paralysis in Aβ transgenic worms (Fig 5A) with the maximum delay, from 24 hours to 72 hours, occurring at 10μM TMG (p<0.0001). 10μM TMG was then used in subsequent experiments. Suppression of *oga-1* by RNAi also delayed Aβmediated paralysis from 24 hours to 48 hours (p<0.0001). In contrast, no change in the median time to paralysis occurred following *ogt-1* gene suppression using RNAi (Fig 5B).

**Figure 5:**
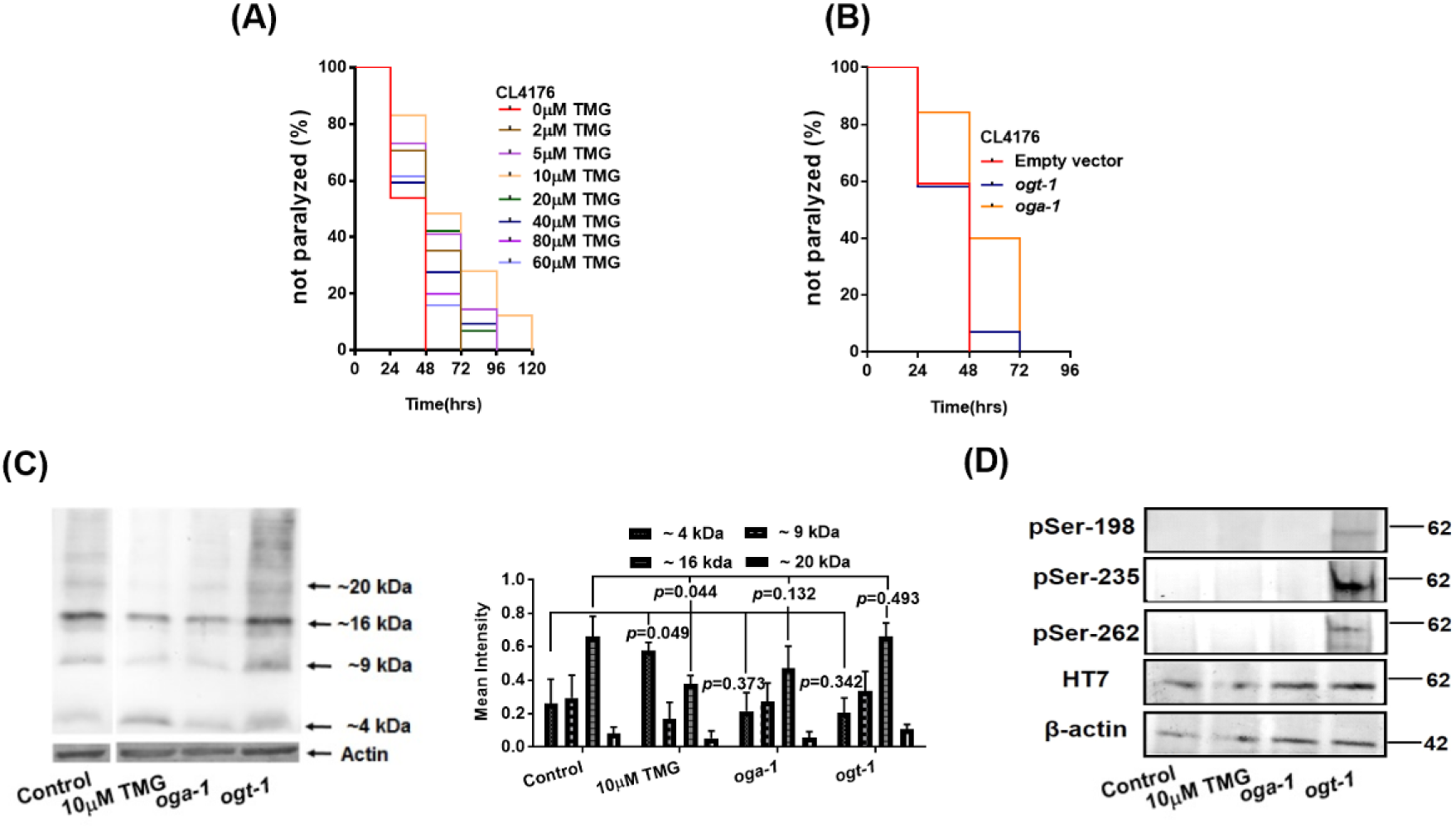
Effect of *oga-1* or *ogt-1* suppression on Aβ-mediated paralysis and tau phosphorylation. *Oga-1* was suppressed either by using Thiamet-G (TMG) or RNAi while *ogt-1* suppression was done using specific RNAi. Temperature inducible Aβ worm strain CL4176 and human fetal tau expressing strain VH255 were used in these experiments. A) paralysis profile of worms that express Aβ fed with different concentrations of TMG ranging from 0-80μM in standard NGM plates. B) Paralysis profile of worms that express Aβ either fed with *ogt-1* or *oga-1* RNAi. Three independent trials were run for each group (n=50-60 worms per trial), and results were presented as the weighted average. Log-rank test was used to determine any significant change in paralysis behavior between each group. C) Effect of 10μM TMG and *oga-1*, and *ogt-1* suppression on Aβ oligomerization in *C. elegans.* D) Effect of 10μM TMG and *oga-1* or *ogt-1* RNAi suppression on site-specific phosphorylation in tau expressing *C. elegans.* After synchronization, VH255 worms were placed at NGM plates containing 10μM TMG or specific *oga-1* or *ogt-1* RNAi. Total cell lysate protein was extracted from 3 days old worms. Tau phosphorylation was monitored using different phosphor-tau-antibodies while total tau protein expression was measured using HT7 antibody. Anti-actin antibody was used as reference control.

The addition of 10μM TMG to the growth medium significantly reduced the levels of Aβ oligomers (~16kDa, 1.75-fold) with a concomitant increase in Aβ monomers (~4kDa, 2.20-fold) compared to untreated control worms (Fig 5C). RNAi mediated suppression of *oga-1* or *ogt-1* gene does not affect the Aβ oligomerization pattern in worms. These results show that suppression of *oga-1* by TMG has a different mechanistic effect than RNAi suppression.

We also determined whether O-β-GlcNAcylation can alter tau phosphorylation by modulating O-β-GlcNAcylation levels. In 3 day old worms, suppression of OGA-1 activity, either by TMG or *oga-1* RNAi, has no effect on tau phosphorylation (Fig 5D). However, *ogt-1* suppression by RNAi significantly increased phosphorylation on Ser198, Ser235 and Ser262 without affecting total tau (HT7). We found no significant change in tau phosphorylation in control worms. Our results indicate that decreasing O-GlcNAcylation resulted in increased tau phosphorylation. At 7 days of age, tau was phosphorylated in control worms, but this was significantly reduced by suppression of the *oga-1* gene with RNAi or suppression of the enzyme activity with TMG (Fig S3).

### Glucose induces Aβ oligomerization and tau phosphorylation in the presence of TMG

Given that glucose increases and TMG decreases Aβ oligomerization, we next investigated the effect of co-treatment. It is evident from Fig 6A that in worms that express Aβ, TMG alone results in lower levels of 9kDa and 16 kDa oligomers than in any of the worms that are exposed to glucose. Exposure to glucose, either with or without TMG caused an increase in Aβ oligomerization. TMG had no protective effect.

**Figure 6:**
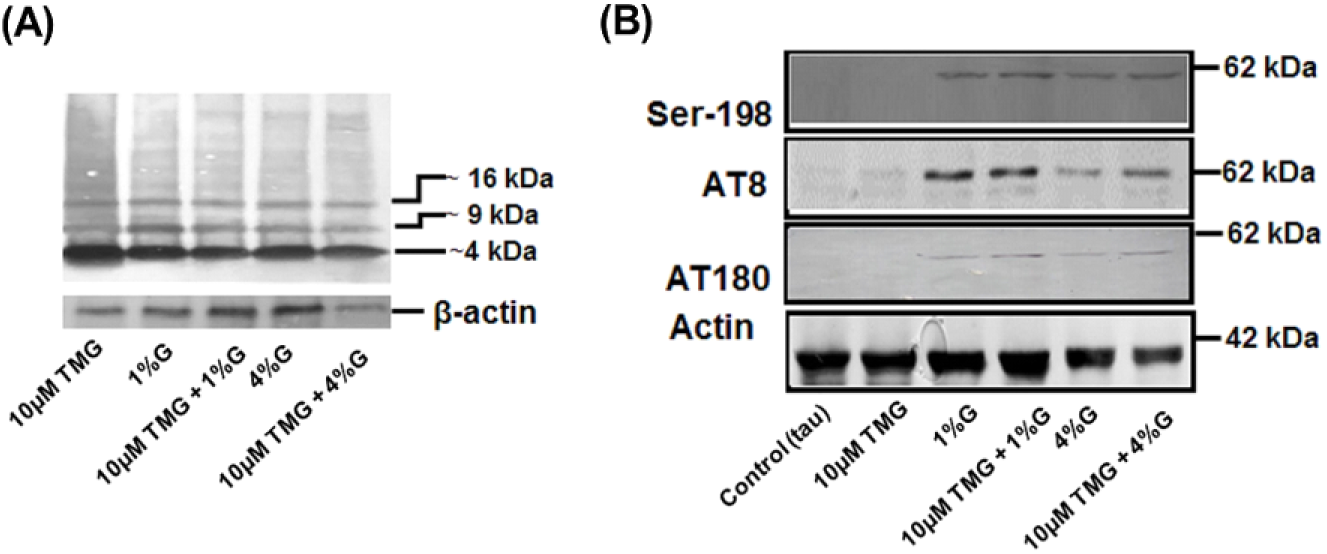
Effect of glucose on Aβ oligomerization, tau phosphorylation in the presence of 10μ? TMG. Synchronized L1 worms expressing either Aβ (CL4176) or tau (VH255) were fed on standard NGM plates containing no glucose or TMG, with 10μM TMG, or 1% glucose or 4% glucose or a combination of 10μM TMG with 1% glucose or 10μM TMG with 4% glucose. Protein was extracted after 72 hrs. A) Effect on Aβ oligomerization. B) Tau phosphorylation was detected on different epitopes from 3 days old worms using conventional SDS-PAGE.

**Fig 7:**
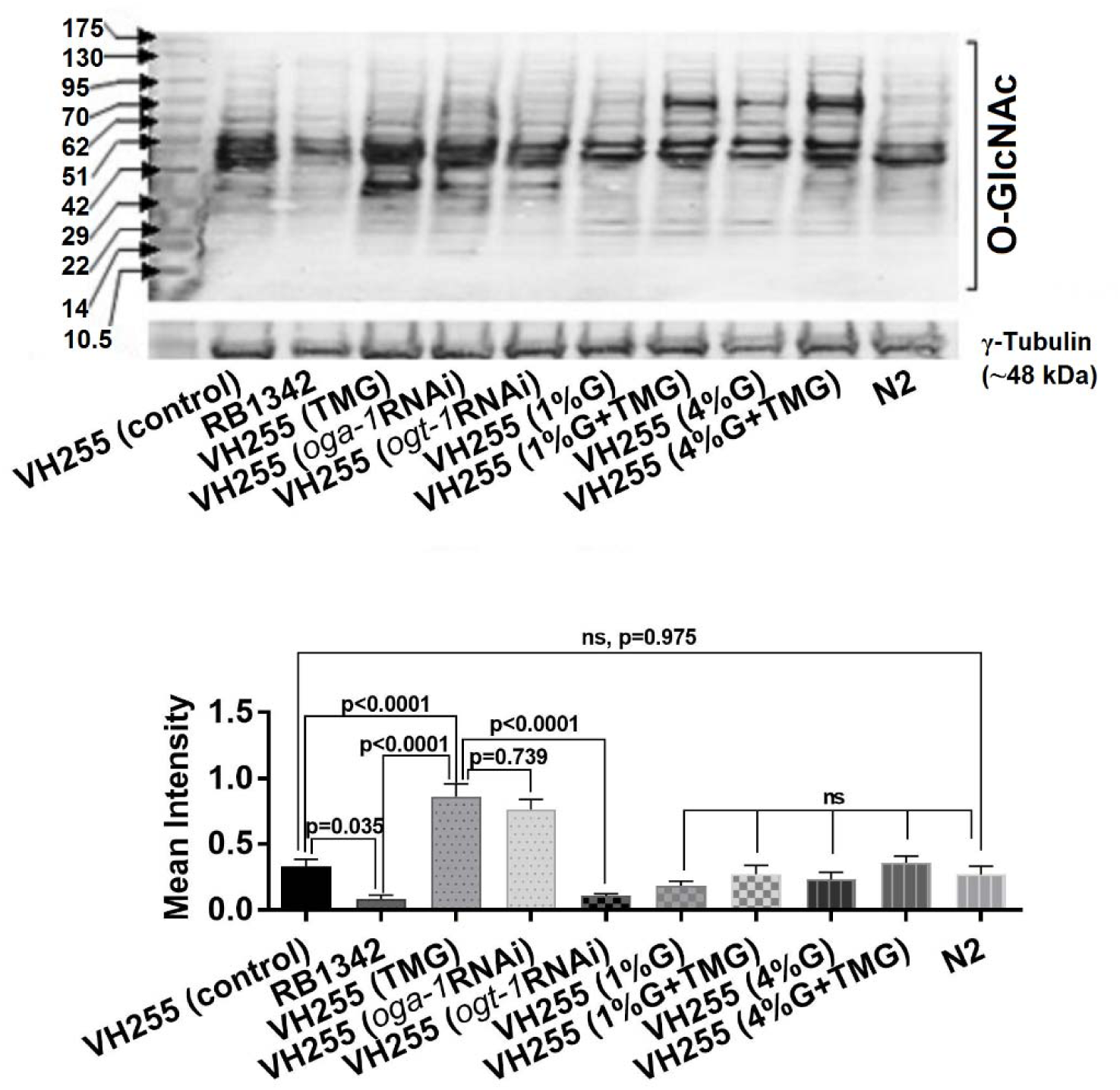
Effect of TMG and glucose on global O-β-GlcNAcylation levels in the worms. Synchronized L1 stage worms were fed on standard NGM plates containing no glucose or TMG, with 10μM TMG, or 1% glucose or 4% glucose or a combination of 10μM TMG with 1% glucose or 10μM TMG with 4% glucose. We also examine the effect of *oga-1* and *ogt-1* suppression by RNAi on O-β-GlcNAcylation. Tau transgenic worm strain VH255 fed with OP50 was presented as the tau control. RB1342 (*ogt-1* deletion) and wild type worms (N2) were run parallel as *ogt-1* −ve control and tau −ve control, respectively. Gel band intensities were quantified using GelQuant.NET. The graphs represent results from two independent trials. Error bars = mean ±SD.

In worms that express tau, only slight phosphorylation of the tau epitopes critical in AD, AT8 and AT180, was observed in the absence of glucose. The level of phosphorylation was not affected by the presence of TMG (AT8, p=0.461; AT180, p=0.251). Glucose alone or in the presence of TMG considerably increased phosphorylation at AT8 (1% G, 7.50 fold, p=0.024; 1%G + TMG, 10.82 fold, p=0.014; 4% G, 10.01 fold, p=0.014; 4%G + TMG, 10.37 fold, p=0.017, respectively) and AT180 1% G (4.81 fold, p=0.043; 1%G + TMG, 5.77 fold, p=0.032; 4% G, 56 fold, p=0.021; 4%G + TMG, 95 fold, p=0.017, respectively) (Fig 6B). As with Aβ oligomerization, glucose cause significant phosphorylation of tau and TMG had no protective effect.

### TMG increases global O-β-GlcNAcylation, but glucose does not

The ability of glucose to induce phosphorylation of tau contradicts the hypothesis that glucose should increase the level of O-β-GlcNAcylation, thereby inhibiting protein phosphorylation. To investigate this further, we monitored the effect of glucose with and without TMG on tau phosphorylation, using TMG, or RNAi of the genes *oga-1* and *ogt-1* as controls. To check whether TMG and glucose are capable of inducing O-β-GlcNAcylation levels in our study, we fed the tau expressing strain VH255 with 10μM TMG and/ or 1% or 4% glucose. We used an *ogt-1* deletion strain RB1342 and wild type N2 as *ogt-1* −ve and tau −ve controls, respectively. We measured the global change in O-β-GlcNAcylation levels rather than individual protein levels. Our results showed that compared to the transgenic tau control VH255, addition of TMG or suppression of *oga-1* increased O-β -GlcNAcylation levels (0.331±0.052 vs 0.864±0.09, 2.61-fold or 0.331±0.052 vs 0.761±0.08, 2.29-fold), Suppression of the *ogt-1* gene by RNAi or an *ogt-1* deletion mutant reduced overall O-β-GlcNAcylation levels relative to the transgenic control strain (0.331±0.052 vs 0.086±0.02, 3.93-fold and 0.761±0.08, 3.04-fold). There was no statistical difference between transgenic and wild type controls (0.331±0.052 vs 0.268±0.06, p=0.975), or between the transgenic control and worms fed with 1% glucose (0.331±0.052 vs 0.184±0.03, p=0.086) or 4% glucose (0.331±0.052 vs 0.232±0.05, p=0.207). Furthermore, use of TMG on glucose fed transgenic worms did not influence O-glycosylation relative to the VH255 transgenic control; 1% or 4% glucose in addition of TMG (0.331±0.052 vs 0.0.271±0.06, p=0.431, (0.331±0.052 vs 0.359±0.04, p=0.631). These results clearly showed that TMG or *oga-1* suppression increased overall O-β-GlcNAcylation levels, whereas glucose did not. Exposure to elevated glucose, however, prevented TMG from increasing O-glycosylation.

## Discussion

Recent studies have linked lifestyle, i.e. carbohydrate-rich diets, to increased neurodegeneration (48, 49). In contrast, lower levels of glucose observed in AD post-mortem brains have been interpreted as one of the major factors responsible for disease progression. These seemingly contradictory observations require further studies to characterize the functional role of glucose in neurodegeneration. In this study, we investigated the effect of glucose enrichment on Aβ and tau-mediated toxicity in *C. elegans.* We also explored the role of O-GlcNAcylation in glucose-mediated changes to tau phosphorylation as well as on Aβ pathogenicity by inhibiting expression of genes encoding enzymes for the glycosylation or de-glycosylation of proteins.

It is well documented that the transgenic expression of human Aβ in the body wall muscle cells of *C. elegans* results in progressive paralysis (38, 50). We found that either 1% glucose or 1% glycerol alleviated paralysis in Aβ transgenic worms. Tauffenberger et al (51) previously found the same result for glucose in worms and suggested that glucose enrichment protects against Aβ proteotoxicity and neurodegeneration. Our results are consistent with this interpretation for both glucose and glycerol. In contrast, in mammalian systems, both glucose and glycerol induce Aβ toxicity (52-54). Interestingly, we also found that 2DOG administration significantly reduced Aβ-mediated paralysis, which is consistent with results in mammalian cell lines and mouse models (21, 22). Understanding this point of contradiction may provide valuable insight into proteotoxicity resulting from hyperglycemia.

We also studied the effect of glucose and 2DOG on ACh neurotransmission in strains that produce Aβ, as extracellular oligomers of Aβ are known to inhibit the release of ACh into the synaptic cleft, resulting in reduced neurotransmission (55-57). ACh neurotransmission is essential for muscle contraction, but increased ACh levels can result in paralysis due to muscle overexcitation. As a result, Aβ expression can provide resistance to cholinergic agonists such as aldicarb, that increases synaptic levels of ACh, or levamisole, that increases ACh receptor sensitization (58, 59). Aldicarb resistance is used as an indication of pre-synaptic defects whereas resistance against both aldicarb and levamisole is used as an indication of post-synaptic defects (30). The presence of Aβ inhibits cholinergic signaling both pre- and post-synaptically by creating resistance against ACh release and blocking axonal vesicle clusters (60-62); we hypothesized that if glucose enrichment can reduce Aβ toxicity, there should be an increase in the sensitivity of Aβ worms to cholinergic hyper-excitation. Contrary to our expectation, glucose induced resistance against both aldicarb and levamisole not only in worms that express Aβ but also in wild type. Thus, glucose enrichment is capable of cholinergic inhibition independent of any direct effect it may have on Aβ.

2DOG induced a very modest level of resistance against both aldicarb and levamisole in wild type worms. In vertebrate models, 2DOG administration induced a stress response in neuronal cells that increased resistance against Aβ and other toxins without changing the normal ACh neurotransmission (63, 64). The partial resistance against aldicarb and levamisole in the presence of 2DOG could be due to the activation of stress resistance proteins as the induced stress response was found to reduces the ACh neurotransmission to compensate for neuronal over-excitation (65). The major observation, however, is that while 2DOG is unable to increase the toxicity of aldicarb in worms that express Aβ, it does restore normal levamisole sensitivity.

Any impairment in ACh signaling could have broad behavioral consequences, e.g. on locomotion, egg laying, mating and muscle contraction (66). Our observation that glucose does not restore cholinergic neurotransmission related to muscle function indicates that it may likewise fail to restore functions impaired by neuronal expression of Aβ. Machino et al. found lower fecundity and increased egg hatching time in worms that express Aβ in neurons. (50). We find that both glucose and glycerol induced further dose-dependent impairment of fecundity and egg viability in worms that express Aβ in neurons. We did not test the effect on wildtype worms, but inhibition of fecundity and egg viability was also seen in *D. melanogaster* and almond moth fed with glucose or glycerol (67, 68). Unlike glucose, exposure to 2DOG did not exacerbate the effect of Aβ on fecundity. Overall, our results strongly demonstrate the negative role of glucose and glycerol on cholinergic neurotransmission.

To help resolve the ambiguous effects of glucose in our study, we examined the effect of a glucose-enriched diet on Aβ oligomerization. Previous studies in mouse models found that hyperglycemia increased Aβ oligomerization (52, 69, 70). As dietary glucose enrichment in our study was reflected in increased intracellular glucose levels in *C. elegans*, it is possible that glucose addition may affect Aβ oligomerization in transgenic worms as well.

Aβ monomers are not toxic, rather the accumulation of Aβ oligomers is responsible for Aβ-mediated neurotoxicity. This is supported by the observation that early cognitive decline coincides with Aβ oligomerization prior to Aβ plaque formation in patients with AD (39, 41, 55-57, 71-76). In our study, glucose gradually induced Aβ oligomerization accompanied by a decrease in Aβ monomers. Our results are consistent with results in mammals, in which carbohydrate-rich diets accelerated the neurodegeneration by increasing Aβ oligomer formation (77, 78). Furthermore, reduction in Aβ oligomerization in mammals resulted in a decrease in disease pathology (34, 38, 59). In our study, we also found that administration of 2DOG reduced cellular dysfunction by decreasing Aβ oligomerization. Similar observation was made in a female mouse model of AD and in cultured neuronal cells, where 2DOG protected against Aβ toxicity by inducing ketogenesis – an alternative energy source used by neurons during glucose deprivation (21, 22).

To further elucidate the role of glucose on AD progression, we assessed the effect of glucose enrichment on tau phosphorylation in tau expressing *C. elegans.* We observed induced phosphorylation on critical tau residues Ser198, Ser235, Ser262, or within clusters of residues AT8 (Ser199/ Ser202/ Thr205) and AT180 (Thr231/ Ser235) after glucose enrichment. The phosphorylation sites examined in our study cause conformational changes in tau and correlate with amino acid residues previously implicated in the severity of neuronal cytopathology in AD (5, 42, 79, 80). Quantitative and kinetic studies have revealed that Ser198/Ser199, Ser202, Thr205, Thr231 and Ser235 are the most critical sites in tau pathology. Phosphorylation at these sites inhibit tau binding with microtubules, resulting in further phosphorylation of tau and promoting self-aggregation (5). Increased tau phosphorylation was also found in mouse models of diabetes when they were injected with streptozotocin, which induces insulin dysfunction and hyperglycaemia suggesting the negative role of glucose on tau phosphorylation (81-83). Our results suggest that glucose enrichment in *C. elegans* might result in insulin-resistance like pathology as proposed in the past (84).

Modulation of tau phosphorylation could be occurred due to the induced O-β-GlcNAcylation that negatively regulate phosphorylation. As glucose, has been demonstrated to promote the O-β-GlcNAcylation of proteins (9, 10), our results indicate the opposite role of glucose in regulating Aβ oligomerization and tau phosphorylation and we hypothesize that glucose enrichment might not induce O-β-GlcNAcylation. This hypothesis was further strengthened by our observation that glucose enrichment did not affect the global O-β -GlcNAcylation in our study. Here we proposed that improving O-GlcNAcylation without disturbing normal glucose levels could be beneficial.

For instance, tau expression was found to reduce worm’s thrashing rates (27), no improvement in thrashing rates was observed in worms in our study either fed with glucose/ glycerol/ 2DOG or TMG or after suppressing *oga-1* and *ogta-1*(Fig S4). These results indicate that reduced thrashing is exclusively associated with tau expression not tau phosphorylation. Similar observations were recorded in the tau expressing *C. elegans* model with pseudohyperphosphorylation, where low or high tau phosphorylation does not affect the thrashing rates (27).

In our study, increased glucose levels in the growth medium resulted in increased levels of glucose in *C. elegans* which is in agreement with an earlier study (84). Importantly, in the current and previous studies using *C. elegans* Aβ models; paralysis has been used as the primary indicator of Aβ toxicity. The most probable reason glucose enrichment delayed paralysis is that glucose also stimulates accelerated movement in the worm. Glucose also keeps the worms alive even when they are continuously experiencing Aβ-toxicity. High glucose administration for longer periods may lead to inhibition of glyoxalase-1 system and production of excess ROS and may enhance disease severity via insulin resistance (47, 48). Our results showed that rather than relying exclusively on paralysis as an indicator of Aβ pathology in *C. elegans*, it is important to use other tools. Collectively, our results suggest that glucose enrichment might not be a suitable therapy to inhibit the progression of AD. However, induction of O-β-GlcNAcylation without increasing the normal glucose level could lead to neuroprotection.

---

**Supplementary information Figure S1**

**Figure S1:**
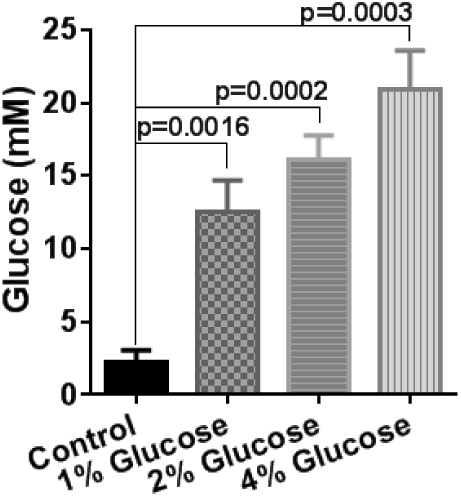
Whole-body glucose levels increased after growth on medium supplemented with glucose. Glucose levels in *C. elegans.* Synchronized wild type *C. elegans* were cultured on NGM plates having the concentrations of 0%, 1%, 2%, and 4% glucose. Concentration of glucose was determined at day 3 in whole-body extracts. Final concentrations were normalized by whole-body protein levels and presented as mM (1% glucose = 55 mM) To determine whether the presence of glucose in NGM plates affects intracellular glucose concentrations of *C. elegans*, we cultured *C. elegans* on different percentage concentrations of glucose in NGM media. We found that a gradual increase in external concentration of glucose also induced intracellular levels of glucose in *C. elegans.* At 0% (control), 1%, 2% and 4% of glucose concentration in NGM resulted in intracellular glucose concentration of 2.21±0.82mM, 12.47±2.19mM, 109±1.67mM, and 20.86±2.69 mM, respectively. These results showed that increases in external glucose levels also increase intracellular glucose concentration in *C. elegans*.

**Figure S2:**
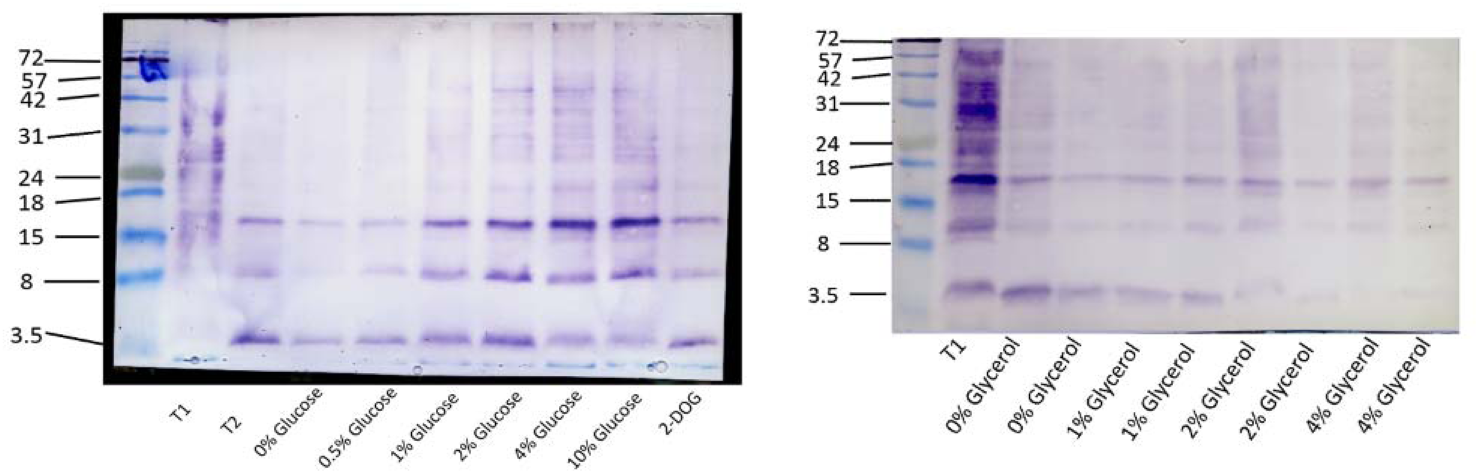
Uncropped images of western blots presented in figure 3. Prism ultra-protein ladder (ab116029) was used to estimate protein molecular weights.

**Figure S3:**
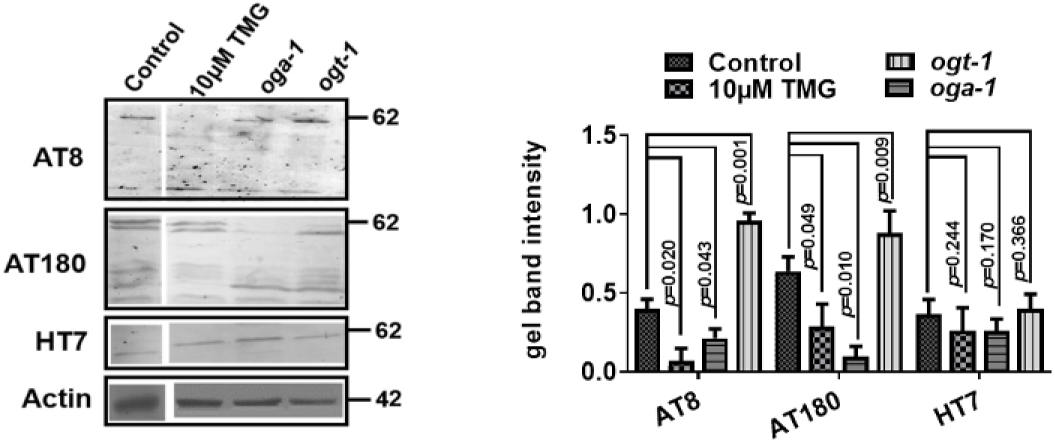
Effect of 10μM TMG and *oga-1* or *ogt-1* RNAi suppression on site-specific phosphorylation in tau expressing *C. elegans.* After synchronization, VH255 worms were placed at NGM plates containing 10μ? TMG or specific *oga-1* or *ogt-1* RNAi. Total cell lysate protein was extracted from 7 day old worms. Tau phosphorylation was monitored using different phosphor-tau-antibodies while total tau protein expression was measured using HT7 antibody. Anti-actin antibody was used as reference control. The graphs represent results from two independent trials. Error bars = mean ±SD. In 7 day aged worms, phosphorylation was apparent in both control and experimental groups. *Oga-1* suppression either by TMG or *oga-1* RNAi knockdown reduced phosphorylation levels at AT8 (5.62 fold, p=0.02; 1.87 fold, p=0.043, respectively) and AT180 (2.21 fold, p=0.049, 52 fold, p=0.01, respectively) in 7 days old worms. Meanwhile *ogt-1* knockdown induced phosphorylation at AT8 2.38 fold, p=0.001) and AT180 (1.37 fold, p=0.09) when compared to control. No change in expression of HT7 levels endorses the equal loading of tau protein that was further confirmed by anti-actin antibody. Our results indicate that with *oga-1* suppression, phosphorylation at tau phosphor-enable residues could be modulated.

**Figure S4:**
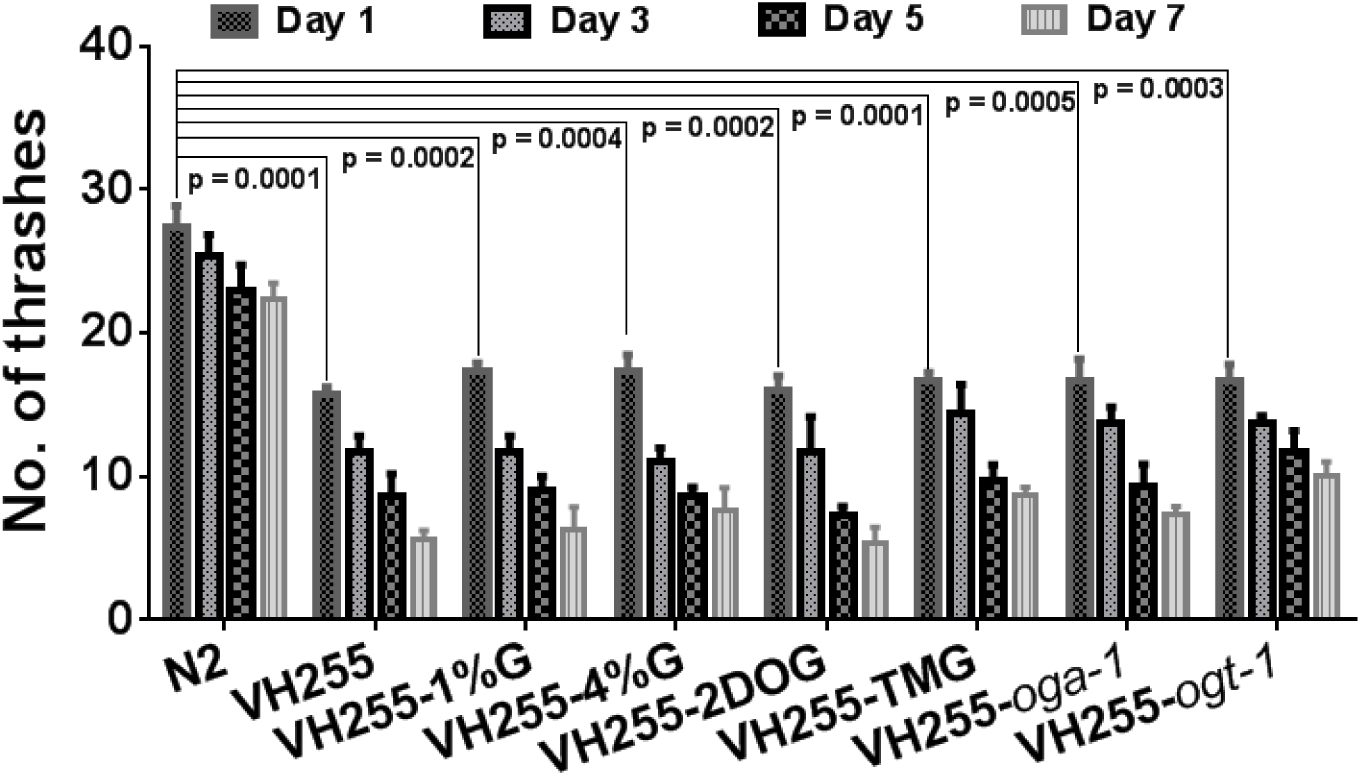
from three independentThrashing assay of wild type and tau expressing worms either fed with glucose, 10μ? TMG or with *oga-1* and *ogt-1* knockdown. Thrashing rates were counted every 2^nd^ day till day 7. After day 3, worms were shifted to the new NGM plates with 7μM FUDR to block the progeny production. We selected 10 worms randomly from each trial and subjected to count for thrashing in M9 buffer for 10sec at 20°C. Neither treatment improve thrashing rates of tau expressing worms. Results were generated from three independent trials. Error bars = mean ± SD. Expression of human tau reduces the thrashing rate in transgenic animals (27). We assessed whether glucose, 2DOG, TMG treatment, and/or *oga-1* or *ogt-1* knockdown affect the trashing rates of tau expressing worms. Tau transgenic worms were assessed for thrashing rates with or without presence of 1% and 4% glucose, or 5mM 2DOG. A gradual decrease in worms thrashing rates was found to be independent of the mode of treatment. We observed similar results for worms fed with TMG or with *oga-1* RNAi or *ogt-1* RNAi.

**Figure S5:**
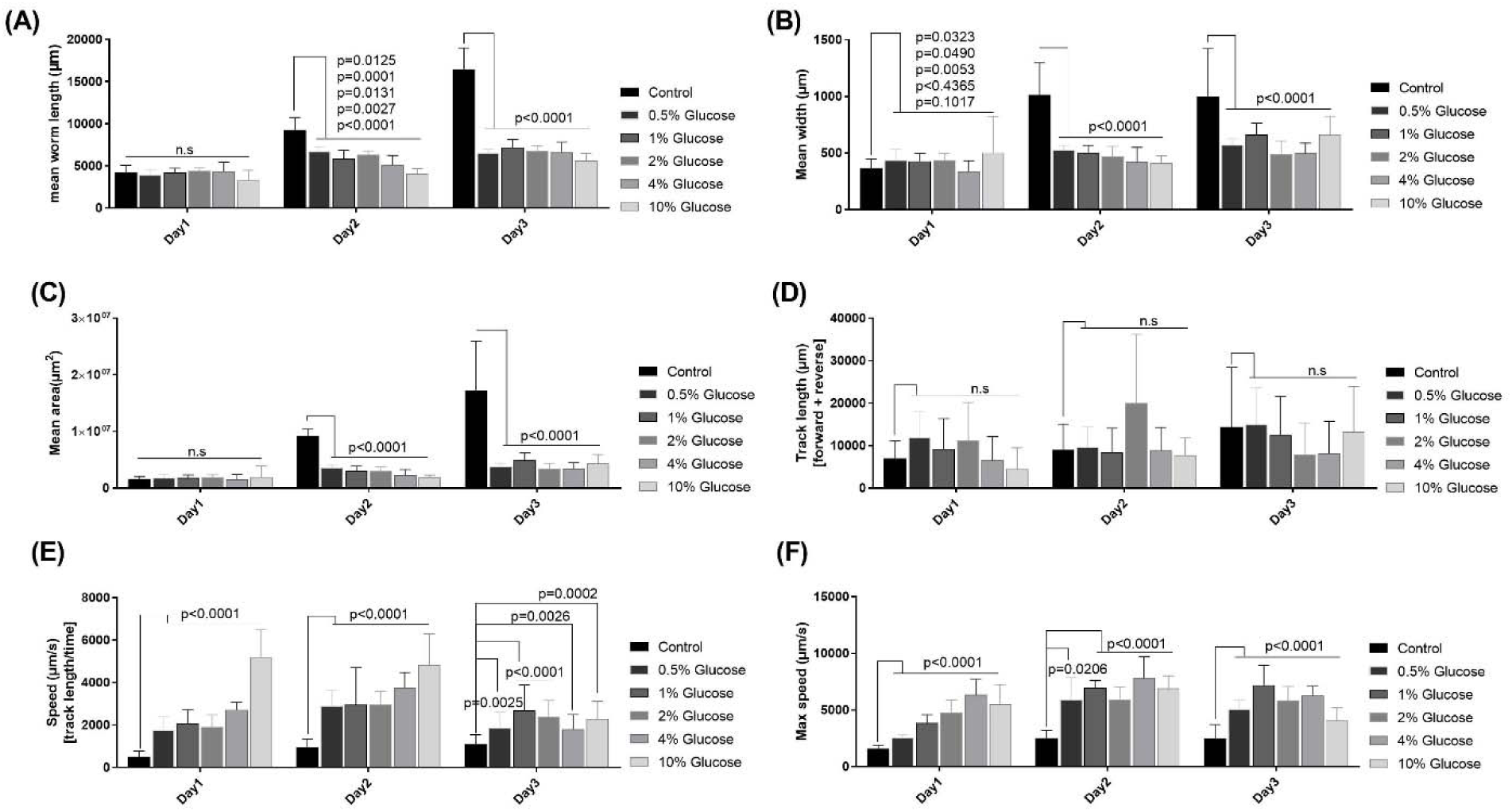
Effect of glucose on body size and speed of the worms. Synchronized wild type worms were treated with different concentrations of glucose on NGM plates seeded with OP50. Videos (30 sec) were recorded and imported to WormLab 4.0 analytical software after 24 hours. We measured the (A) worm’s mean length, (B) width, (C) area, (D) track length, (E) average speed and (F) maximum speed. At least 10 worms were selected randomly from each cohort. Error bars = mean±SD. We observed increased movement of worms that express Aβ in muscle despite induced Aβ oligomerization when treated with glucose. These results were unexpected and we further tested the effect of glucose on body size and movement of the wild type worms. Addition of glucose significantly induced the speed of worms. However, after 48 hours, both average and maximum speed was starts to drop for worms treated with 4% or 10% glucose suggesting the toxic effects of glucose at high concentrations. Overall, our results emphasis impaired body shape and speed of worms after treatment with glucose.

**Table S1:** Statistical comp arison of paralysis curves after exposure to glucose or 2DOG in the presence of aldicarb or levamisole

|  |  |  |  |  |
| --- | --- | --- | --- | --- |
| **Treatment** | **N2 (aldicarb)** |  | **N2 (levamisole)** |  |
|  | Median time to survival (min) | P value (log rank) | Median time to survival (min) | P value (log rank) |
| 0% glucose | 30 |  | 30 |  |
| 0.5% glucose | 30 | 0.0004 | 90 | <0.0001 |
| 1% glucose | 60 | <0.0001 | 90 | <0.0001 |
| 2% glucose | 60 | <0.0001 | 90 | <0.0001 |
| 4% glucose | 60 | <0.0001 | 90 | <0.0001 |
| 5mM 2DOG | 60 | <0.0001 | 60 | <0.0001 |
| **Treatment** | **CL2006 (aldicarb)** |  | **CL2006 (levamisole)** |  |
|  | Median survival (min) | P value (log rank) | Median survival (min) | P value (log rank) |
| 0% glucose | 60 |  | 60 |  |
| 0.5% glucose | 60 | 0.1074 | 90 | 0.043 |
| 1% glucose | 90 | 0.0011 | 90 | 0.009 |
| 2% glucose | 90 | <0.0001 | 90 | 0.001 |
| 4% glucose | 90 | <0.0001 | 90 | <0.0001 |
| 5mM 2DOG | 60 | 0.589 | 30 | <0.0001 |

